# RFPlasmid: Predicting plasmid sequences from short read assembly data using machine learning

**DOI:** 10.1101/2020.07.31.230631

**Authors:** Linda van der Graaf van Bloois, Jaap A. Wagenaar, Aldert L. Zomer

**Affiliations:** Faculty of Veterinary Medicine, Department of Infectious Diseases and Immunology, Utrecht University, Utrecht, the Netherlands; WHO Collaborating Centre for Reference and Research on Campylobacter and Antimicrobial Resistance from an One Health Perspective/ OIE Reference Laboratory for Campylobacteriosis, Utrecht, the Netherlands; Wageningen Bioveterinary Research, Lelystad, the Netherlands

## Abstract

Antimicrobial resistance (AMR) genes in bacteria are often carried on plasmids and these plasmids can transfer AMR genes between bacteria. For molecular epidemiology purposes and risk assessment, it is important to know if the genes are located on highly transferable plasmids or in the more stable chromosomes. However, draft whole genome sequences are fragmented, making it difficult to discriminate plasmid and chromosomal contigs. Current methods that predict plasmid sequences from draft genome sequences rely on single features, like k-mer composition, circularity of the DNA molecule, copy number or sequence identity to plasmid replication genes, all of which have their drawbacks, especially when faced with large single copy plasmids, which often carry resistance genes. With our newly developed prediction tool RFPlasmid, we use a combination of multiple features, including k-mer composition and databases with plasmid and chromosomal marker proteins, to predict if the likely source of a contig is plasmid or chromosomal. The tool RFPlasmid supports models for 17 different bacterial species, including *Campylobacter*, *E. coli*, and *Salmonella*, and has a species agnostic model for metagenomic assemblies or unsupported organisms. RFPlasmid is available both as standalone tool and via web interface.

## Introduction

Many bacterial species carry plasmids, extrachromosomal mobile genetic elements that can transfer from one bacterium to another (Smillie et al., 2010). They often replicate autonomously in the host using a variety of replication systems. Generally they are circular, however some species carry linear plasmids (Dib et al., 2015; Li et al., 2007). Plasmids carry often genes which provide a benefit to the host, such as additional metabolic capabilities (Rozwandowicz et al., 2018), antimicrobial resistance genes (Carattoli, 2009) and virulence factors that affect host invasion and infection, including type IV secretion systems, toxins, adhesins, invasins and antiphagocytic factors (Johnson & Nolan, 2009; Sengupta & Austin, 2011). The presence of plasmids increases in general the fitness of their hosts by providing new functions, like antimicrobial resistance (AMR), metabolic capabilities or virulence factors.

Conjugative transfer of plasmids is considered the most effective way of spreading antimicrobial resistance among bacteria (Goessweiner-mohr et al., 2014). For molecular epidemiology purposes and risk assessment, the identification of chromosomal and plasmid sequences provides fundamental knowledge regarding the transmission of AMR. Molecular identification of plasmid and chromosomal genotypes can distinguish whether the spread of AMR genes is driven by epidemic plasmids to different hosts or by clonal spread of bacterial organisms.

Many molecular epidemiology studies using short read Illumina sequences are available for resistant organisms and the number of sequenced genomes available is in the hundreds of thousands (Alikhan et al., 2018; Jolley et al., 2018; Wattam et al., 2017). These existing datasets could provide a wealth of information on plasmid dissemination, were it not for one major drawback: assembly of short read sequencing data results in hundreds of contigs which are difficult to circularize conclusively, making it next to impossible to determine what is plasmid and what is chromosomal DNA.

Multiple bioinformatic methods have been described to predict plasmids *in silico*, e.g. cBar (Zhou & Xu, 2010), PlasmidSPAdes (Antipov et al., 2016), Recycler (Rozov et al., 2017), PlasmidFinder (Carattoli et al., 2014), PLACNET (Lanza et al., 2014), PLAScope (Royer et al., 2018), MLPlasmids (Arredondo-Alonso et al., 2018) and Platon (Schwengers et al., 2020). The predictions with some methods suffer from a low sensitivity or specificity (Arredondo-Alonso et al., 2017), or are optimized for one specific bacterial genus and cannot be used for metagenomics.

In this study, we present our tool RFPlasmid, a novel approach for the prediction of bacterial plasmid sequences in contigs from short read assemblies, with models for 17 different bacterial genera and a species agnostic model. We compared RFPlasmid to other available tools and show it that it performs equally well or better when using species-specific models. We identified genomic signatures of plasmid and chromosomal sequences based on 5-basepair k-mers, a custom plasmid protein database with >193,000 entries, a database of known replicons (Zankari et al., 2012), single copy chromosomal marker genes (Parks et al., 2015), contig lengths and gene counts. We trained a Random Forest model on more than 8000 pseudo assemblies from bacterial chromosomes and plasmids and validated our approach using both the out of bag (OOB) error rate of Random Forest and an independently generated dataset of plasmid and genomic contigs. Our prediction model achieves up to 99% classification accuracy and 99% sensitivity on genome assemblies of bacterial species of 17 different genera and metagenomics, outperforming any other tool currently available. Additionally, we have identified potential factors responsible for prediction errors and propose downstream analyses to alleviate these problems.

## Implementation

RFPlasmid extracts feature information from whole genome sequences (WGS) contigs and by using a Random Forest model, the likely source (plasmid or chromosomal) of the contigs is predicted. The tool supports 17 different bacterial species or genera, including *Bacillus, Borrelia, Burkholderia, Campylobacter, Clostridium, Corynebacterium, Cyanothece, Enterobacteriaceae, Enterococcus, Lactobacillus, Lactococcus, Listeria, Pseudomonas, Rhizobium, Staphylococcus, Streptomyces, Vibrio* and a species agnostic model for unknown unsupported organisms or for metagenomics data. A flow scheme describing the procedure is given in Figure 1. Furthermore the tool contains an easy to use training option with which additional models can be added easily.

**Figure 1.**
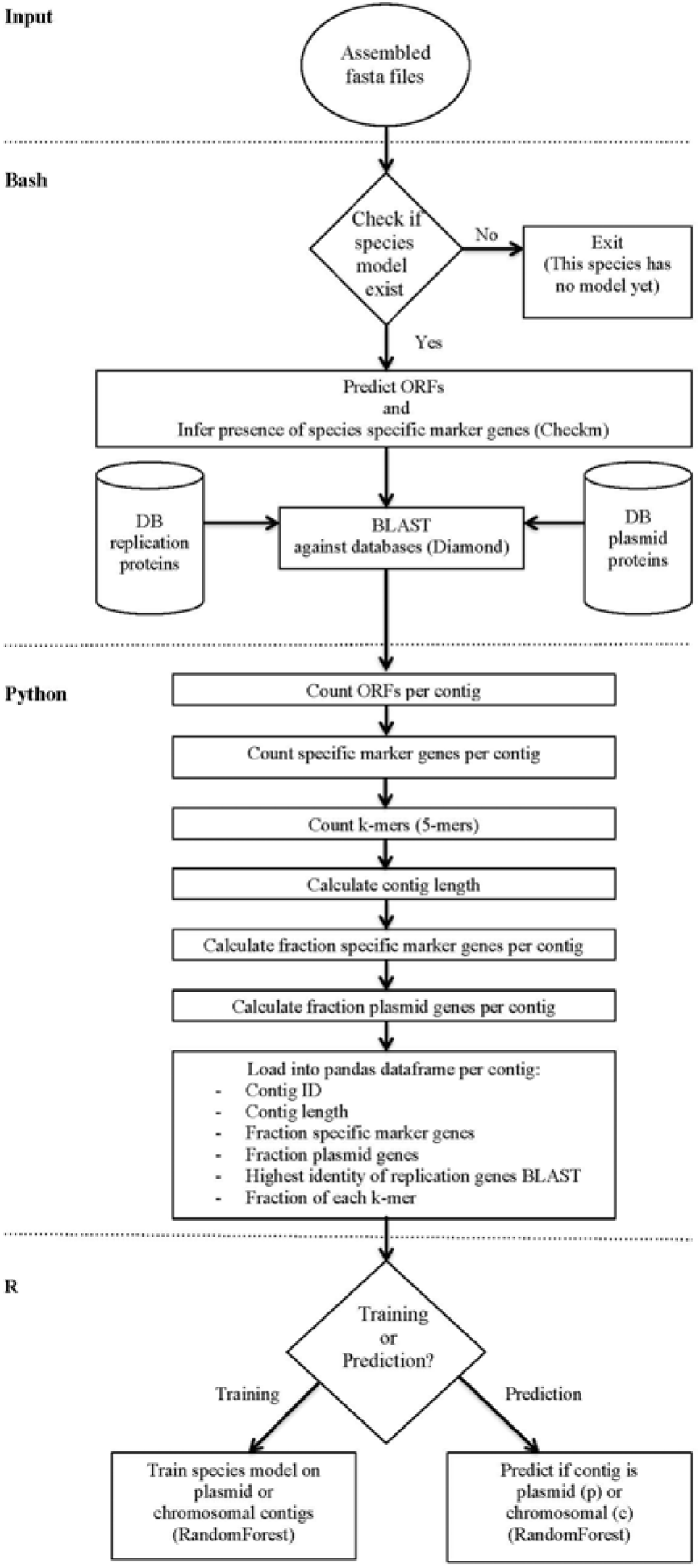
Flow diagram of RFPlasmid.

### Input

Contigs from short read assemblies in FASTA format are used as input files. The webinterface takes a single genome, the command line tool can process up to several thousand genomes from a single folder.

### Single copy chromosomal marker genes

CheckM (Parks et al., 2015) predicts open reading frames (ORFs) of the contigs using Prodigal (Hyatt et al., 2010) and determines whether these encode species specific single¬copy marker genes. The number of specific marker genes per contig is counted and saved.

### Plasmid marker proteins

Two different reference databases with plasmid maker proteins are used; the plasmid replicon database and plasmid protein database. The plasmid replicon database consists of known plasmid replication proteins, downloaded from the database of PlasmidFinder (Zankari et al., 2012) (accession date: 22 May 2017). The plasmid protein database was generated with plasmid proteins from all bacterial species from NCBI Genbank (accession date: 22 May 2017) and the plasmid database of the MOB-suite (Robertson & Nash, 2018). Near-identical proteins were clustered using USearch v 5.2.32 (Edgar, 2010), resulting in a database with 193,176 plasmid proteins.

RFPlasmid uses DIAMOND searches (Buchfink et al., 2015) against the two plasmid reference databases, BLASTX for the replicon database and BLASTP for the protein database, with default settings and an E-value cutoff of 1E-30. For each contig, the BLASTX replicon hit with the highest identity is selected and the number of BLASTP hits with the plasmid protein database is counted.

### K-mer profiles

Two different methods of k-mer counting are implemented; the standard method counting the number of nucleotide pentamers (5-mers) using python (default), and the faster, optional method JELLYFISH (Marçais & Kingsford, 2011). The fraction of each 5-mer is calculated.

### Classifying using Random Forest models

A Python Pandas dataframe is generated, to structure all the different features of the query WGS contigs, including contig name, contig length, fraction specific maker genes, fraction plasmid genes, highest replication gene identity and k-mer fractions. The Pandas dataframe is exported as a csv, which is imported in R for training or classification using the Random Forest library (Liaw & Wiener, 2002).

#### Training data sets

The training data sets were made as follows; complete and identified chromosomal and plasmid sequences were downloaded from NCBI Genbank (accession date 7 November 2017), and for *Listeria*, plasmid sequences were downloaded from NCBI Genbank with accession date 30 September 2019. Pseudo reads of 500bp each were generated with 50x coverage using the gen-single-reads script (https://github.com/merenlab/reads-for-assembly). Assembly was performed using SPAdes v3.11.1 (Bankevich et al., 2012) with default settings. Contigs smaller than 200 bp were removed. Table 1 shows the assemblies of the developed training data sets of each species.

**Table 1.** Assemblies of the developed training data sets.

| Species | Number of chromosomes | Number of Plasmids | Number of generated contigs for training (c – p) | Total number of basepairs |
| --- | --- | --- | --- | --- |
| *Bacillus* | 377 | 291 | 20055<br>(15736 – 4319) | 1.77E+09 |
| *Borrelia* | 28 | 23 | 1564<br>(110 – 1454) | 3.32E+07 |
| *Burkholderia* | 211 | 47 | 26256<br>(25139 – 1118) | 1.48E+09 |
| *Campylobacter* | 197 | 406 | 5423<br>(3652 – 1771) | 3.42E+08 |
| *Clostridium* | 100 | 46 | 6537<br>(6044 – 493) | 4.09E+08 |
| *Corynebacterium* | 166 | 63 | 4614<br>(4350 – 264) | 4.31E+08 |
| *Cyanothece* | 5 | 6 | 634<br>(399 – 235) | 2.95E+07 |
| *Enterobacteriaceae* | 151 | 2297 | 28544<br>(13621 – 14923) | 9.07E+08 |
| *Enterococcus* | 57 | 44 | 6270<br>(3693 – 2576) | 1.73E+08 |
| *Lactobacillus* | 206 | 110 | 19412<br>(15610 – 3802) | 5.17E+08 |
| *Lactococcus* | 37 | 76 | 3423<br>(2104 – 1319) | 9.04E+07 |
| *Listeria* | 142 | 73 | 2685<br>(2371 – 200) | 4.24E+08 |
| *Pseudomonas* | 254 | 42 | 18645<br>(17636 – 1009) | 1.58E+09 |
| *Rhizobium* | 52 | 51 | 4241<br>(1573 – 2668) | 3.50E+08 |
| *Staphylococcus* | 247 | 136 | 9124<br>(7763 – 1361) | 6.81E+08 |
| *Streptomyces* | 82 | 64 | 6449<br>(6357 – 92) | 7.03E+08 |
| *Vibrio* | 123 | 41 | 11265<br>(10282 – 983) | 5.91E+08 |
| Metagenomics / bacteria | 2958 | 3937 | 222723<br>(194597 – 28126) | 1.19E+10 |

Random Forest models were trained using 5000 trees. Class imbalances were solved by making use of the sampsize option, whereby 66% of the smallest class was selected as option in sampsize for both classes when training each tree in the forest to prevent class imbalance errors and error inflation (Janitza & Hornung, 2018). Random Forest uses an internal validation where 66% of the contigs of the training-sets are used for training and 33% are used for testing per tree in the Random Forest. The output of every tree is averaged and results in the OOB (out-of-bag) error which is a minor overestimation of the actual error (Janitza & Hornung, 2018).

#### External validation

To investigate the performance of RFPlasmid on non-simulated datam, we downloaded the Illumina and Nanopore reads of 24 multidrug-resistant *Escherichia coli* genomes from ENA from Bioprojects PRJNA505407 and PRJNA387731 which were also used by Schwengers et al (Schwengers et al., 2020). We performed both hybrid assembly using Unicycler v0.4.9b (Wick et al., 2017) and short read-only assembly with SPAdes (v13.3.0). We could assemble 22 isolates into distinct chromosomal and plasmid contigs using Unicycler. Isolates V232 and V92 were excluded after inspection of the sequence graphs using Bandage (Wick et al., 2015) as chromosomal and plasmid contigs could not be distinguished. Contigs larger than 200 bp from the SPAdes assemblies were aligned against the corresponding complete hybrid assembly using Last (Hamada et al., 2017) and the best scoring hits against plasmid and chromosome contigs were collected. In total, 85 contigs (153 kbases) of the 2832 (110 mbases) contigs in the entire dataset were discarded as they had identical hits on both chromosome and plasmid.

#### Analysis of predicted contigs for encoded features

Presence of phage genes and resistance genes in assembled contigs of the training data were determined by performing a DIAMOND search against the ProPHET phage database (Reis-Cunha et al., 2019) using an E-value cutoff of 1E-10 and the Resfinder database (accessed 01-07-2020) with a cutoff of 90% identity and 60% coverage (identical to the default settings of the online version of Resfinder). Presence of transposase encoding genes was performed by aligning encoded proteins using HMMER3 (http://hmmer.org/) against the transposase database of ISEscan (Xie & Tang, 2017) with an E-value cutoff of 1E-30.

#### Software availability

Software is available at https://github.com/aldertzomer/RFPlasmid, databases containing all plasmid proteins are available at http://klif.uu.nl/download/plasmid_db/ and all training data is available at http://klif.uu.nl/download/plasmid_db/trainingsets2/trainingsfiles_zip. A webinterface for RFPlasmid is available at http://klif.uu.nl/rfplasmid/

## Results

### Classification results on training data

Sizes of in this study generated plasmid contigs of the training data sets vary between 200bp and 4,146,943bp, with species *Bacillus* having the largest contig (data not shown). Between 127 and 11513 plasmid contigs per species are available, with the *Enterobacteriaceae* set having the highest number of plasmid contigs (Table 1). We compared the predicted contig location to the known contig location with plasmid contigs correct classified as plasmid (plasmid correct), chromosomal contigs correct classified as chromosomal (chromosome correct), chromosomal contigs incorrect classified as plasmids (chromosome incorrect) and plasmid contigs incorrect classified as chromosomal (plasmid incorrect). Results are determined in percentages, calculated as basepairs of each predicted contig divided by the total basepairs, because smaller contigs are more difficult to predict, making the number of contigs a poor comparator.

To address potential over-training, we present both OOB (out-of-bag) errors and prediction failures of the complete model. Random Forest uses an internal validation where 66% of the contigs of the training-sets are used for training and 33% are used for testing per tree in the Random Forest. The output of every tree is averaged and results in the OOB error which is a minor overestimation of the actual error (Janitza & Hornung, 2018). Both OOB classification results and the output of the complete model on the training data sets are presented in Figure 2A and 2B. The results show that RFPlasmid can correctly identify the source of the contigs between 95.3% and 100%. Often, the species specific model outperforms the species agnostic model (Fig 2AB).

**Figure 2.**
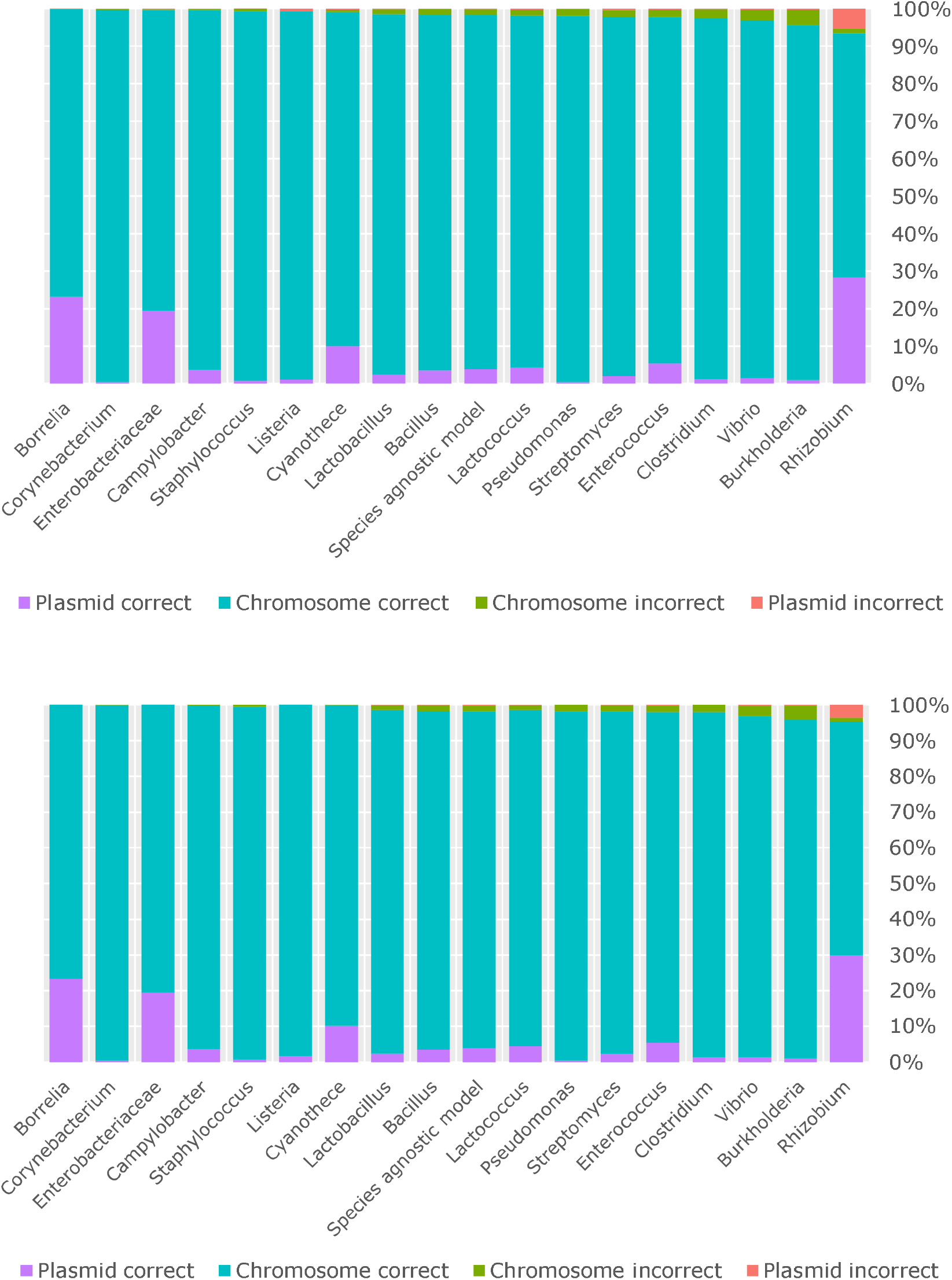
Performance of Random Forest classification models on training data. Shown are (A) OOB performance and (B) prediction performance in percentages, calculated as basepairs predicted divided by the total basepairs for each contig, coloured as plasmid correct, chromosome correct, chromosome incorrect and plasmid incorrect classified contigs.

We observe that contigs that with scores between 0.4 and 0.6 are the main source of incorrectly predicted contigs (Figure 3A). Contigs smaller than 3 kb are difficult to classify, their scores are generally lower (Figure 3B) possibly because the k-mer content cannot be reliably determined, or the contigs do not contain coding sequences (CDS), or they consist of genes that usually have multiple copies on both genome and plasmid such as transposases or phage genes. To investigate the latter hypothesis, we determined the presence of phage genes and transposases on the incorrectly and correctly predicted contigs and determined the phage- and transposas e-content per contig. This analysis was performed on contigs containing at least one CDS. Phage genes were mainly found in chromosome incorrect classified contigs, where 30% (3482 of 11565) of the chromosome incorrect classified contigs harboured phage genes, of which 34% (1179 of 3483) of the contigs consisted of >50% phage genes (Figure 4A). Transposases were found in a very high percentage of 36% (4200 of 11565) of chromosome incorrect classified contigs as well (Figure 4B) and 59% (2487 of 4200) of these transposase carrying contigs were small contigs (<3kb).

**Figure 3.**
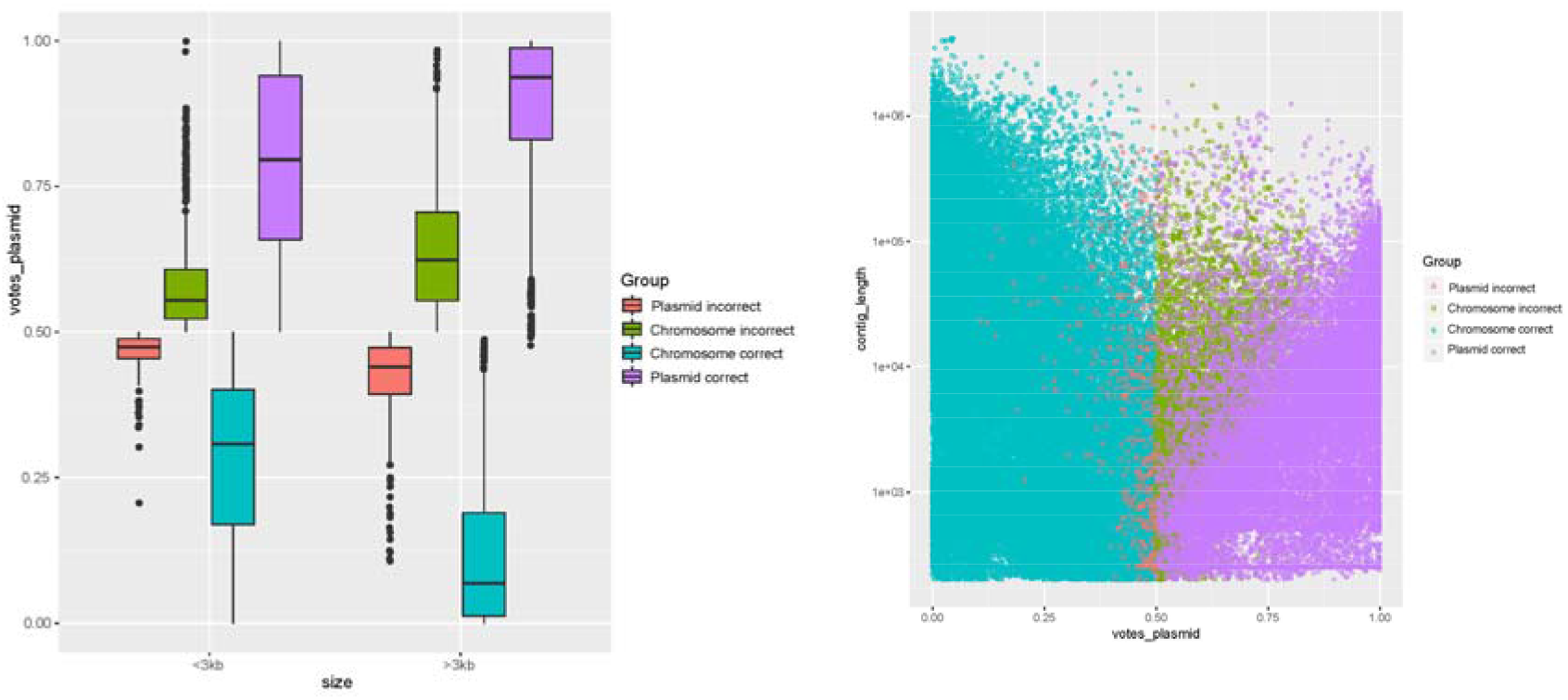
Plots of RFPlasmid prediction scores. (A) Boxplot displaying the plasmid prediction votes of small (< 3 kb) and large (>3 kb) contigs, grouped per correct and incorrect classified plasmid and chromosome contigs. (B) Scatterplot displaying the plasmid prediction votes the contig lengths (log10 scale), coloured by the classification results.

**Figure 4.**
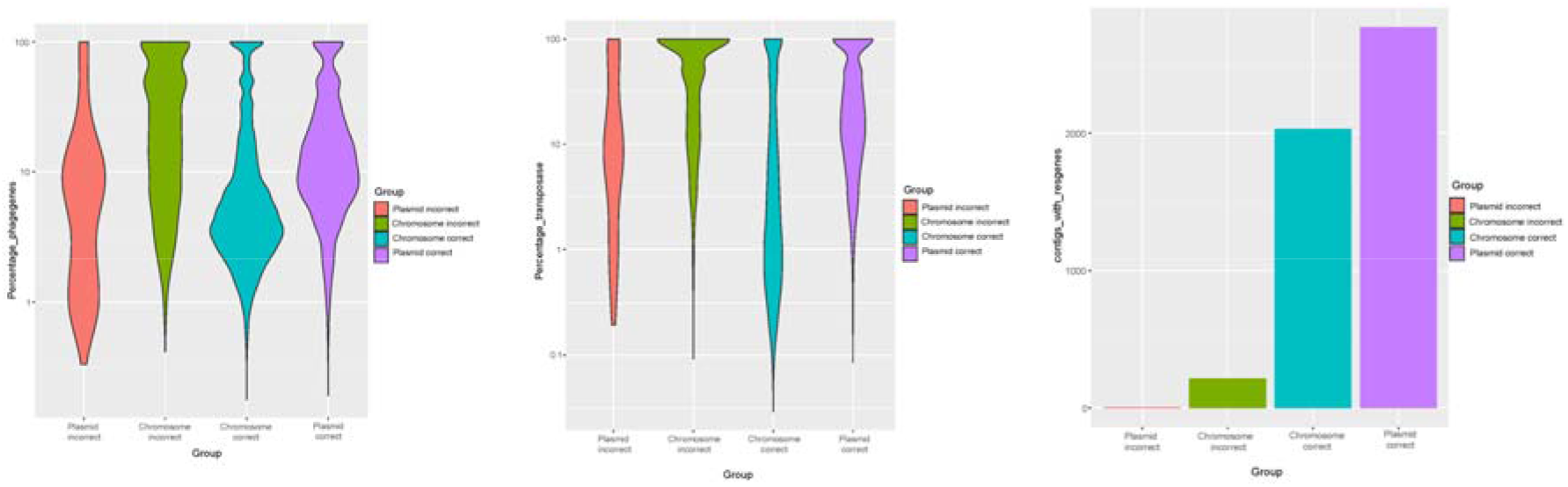
Presence of phage genes, transposases and resistance genes in training data contigs. Shown are (A) the percentage of phage genes (log10 scale) in training data contigs, (B) the percentage of transposases (log10 scale) in training data contigs, and (C) barplot with counts of contigs with ≥ 1 resistance gene, all grouped per correct and incorrect classified plasmid and chromosome contigs.

As the primary reason for our tool is to determine if we can reliably predict whether AMR genes are carried on plasmids or chromosomes, we analysed the assembled contigs for the presence of resistance genes using the Resfinder database. Resistance genes were found on 5019 of the 175027 contigs (135004 contigs with >1 CDS) (Figure 3C), of which 13% (2773 out of 21306) of plasmids contigs carry an AMR gene and 1.77% (1977 out of 112006) of the chromosomal contigs carry AMR genes. Only 0.9% (3 out of 344) of the plasmid incorrect classified contigs contained AMR genes, and 2.2% (213 out of 9464) of the chromosome incorrect classified contigs contained AMR genes. Of these 213 chromosome incorrect classified AMR gene harboring contigs, 38% (n=82) were located on small contigs (<3kb), therefore we conclude that we reliably identify the DNA source that carries these genes for e.g. risk assessment.

Investigating the importance of each feature in the different training models shows that single copy chromosomal markers genes and plasmid marker genes appear to function species wide as they are important in every model, while k-mer content is specific per species (Figure 5).

**Figure 5.**
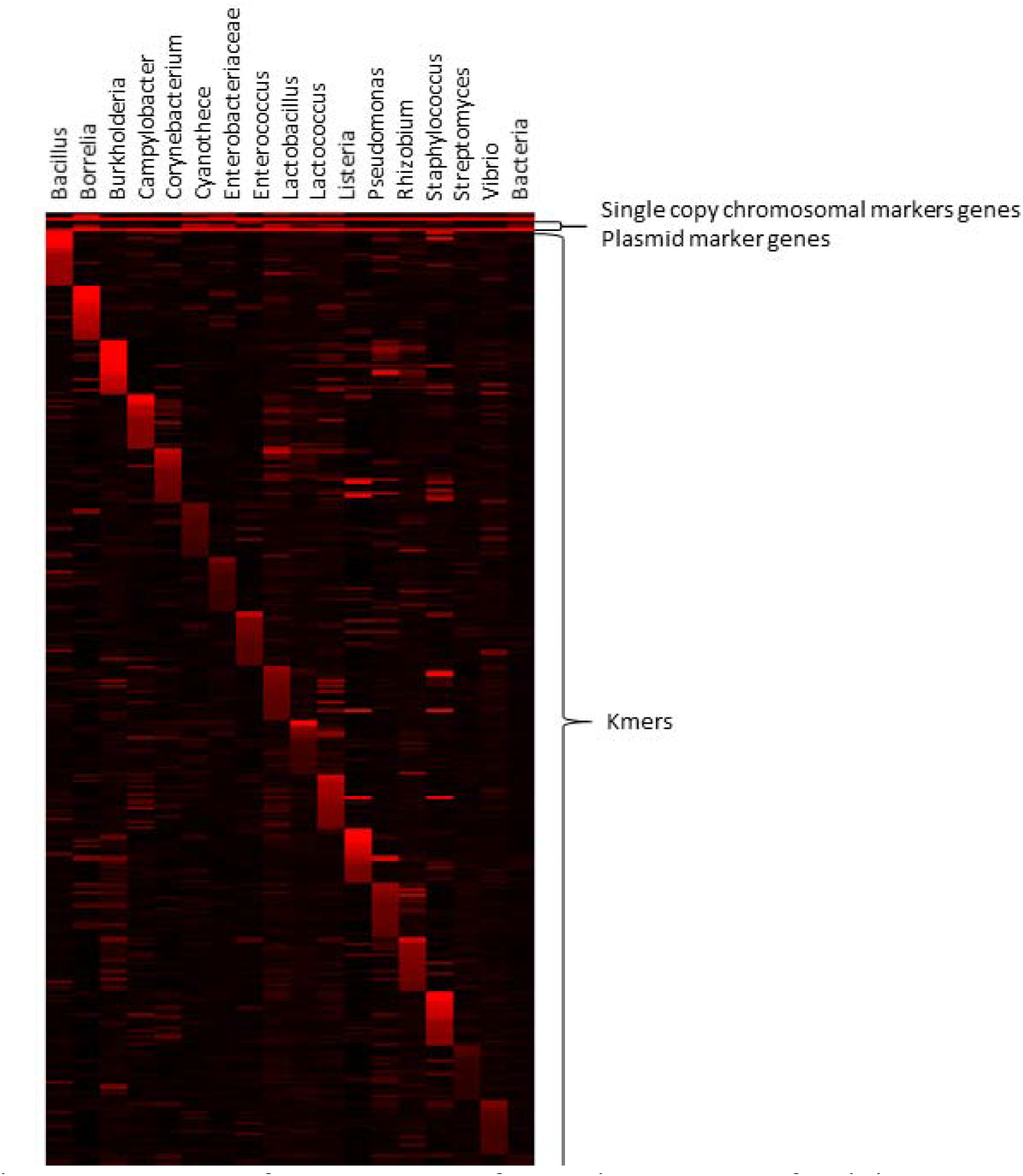
Heatmap of RandomForest feature importances of training models. Features in red were more often selected as discriminatory between chromosomal and plasmid contigs.

### Benchmarking RFplasmid and comparison with existing tools

We compared the performance of RFPlasmid with other plasmid-prediction tools. Plasmid-predictions tools that assemble plasmid contigs from read files like PlasmidSPAdes (Antipov et al., 2016), Recycler (Rozov et al., 2017) and PLACNET (Lanza et al., 2014) are not developed to be used with assembled data, and are therefore excluded in this comparison. The plasmid-prediction tools that can predict plasmid contigs from assembled data were tested and compared with RFPlasmid by using the in this study described models and training data; cBar (Zhou & Xu, 2010) with the metagenome training data, PLAScope (Royer et al., 2018) with the *E. coli* subset of the *Enterobacteriaceae* training data, MLPlasmids (Arredondo-Alonso et al., 2018) with the *Enterococcus* faecium and *E. coli* subsets of the *Enterococcus* training data and *Enterobacteriaceae* training data respectively. Percentages of correctly predicted basepairs are calculated and compared with the RFPlasmid prediction results (Figure 6AB). We show that RFPlasmid outperforms the tested tools by having a lower number of incorrect classified plasmid and chromosome contigs compared to cBar en MLPlasmids for Enterococcus, and by predicting a lower number of plasmid incorrect classified contigs compared to PLAScope. RFPlasmid has an average chromosome incorrect classified contig rate of 1.24% and an average plasmid incorrect classified contig rate of 0.29%.

**Figure 6.**
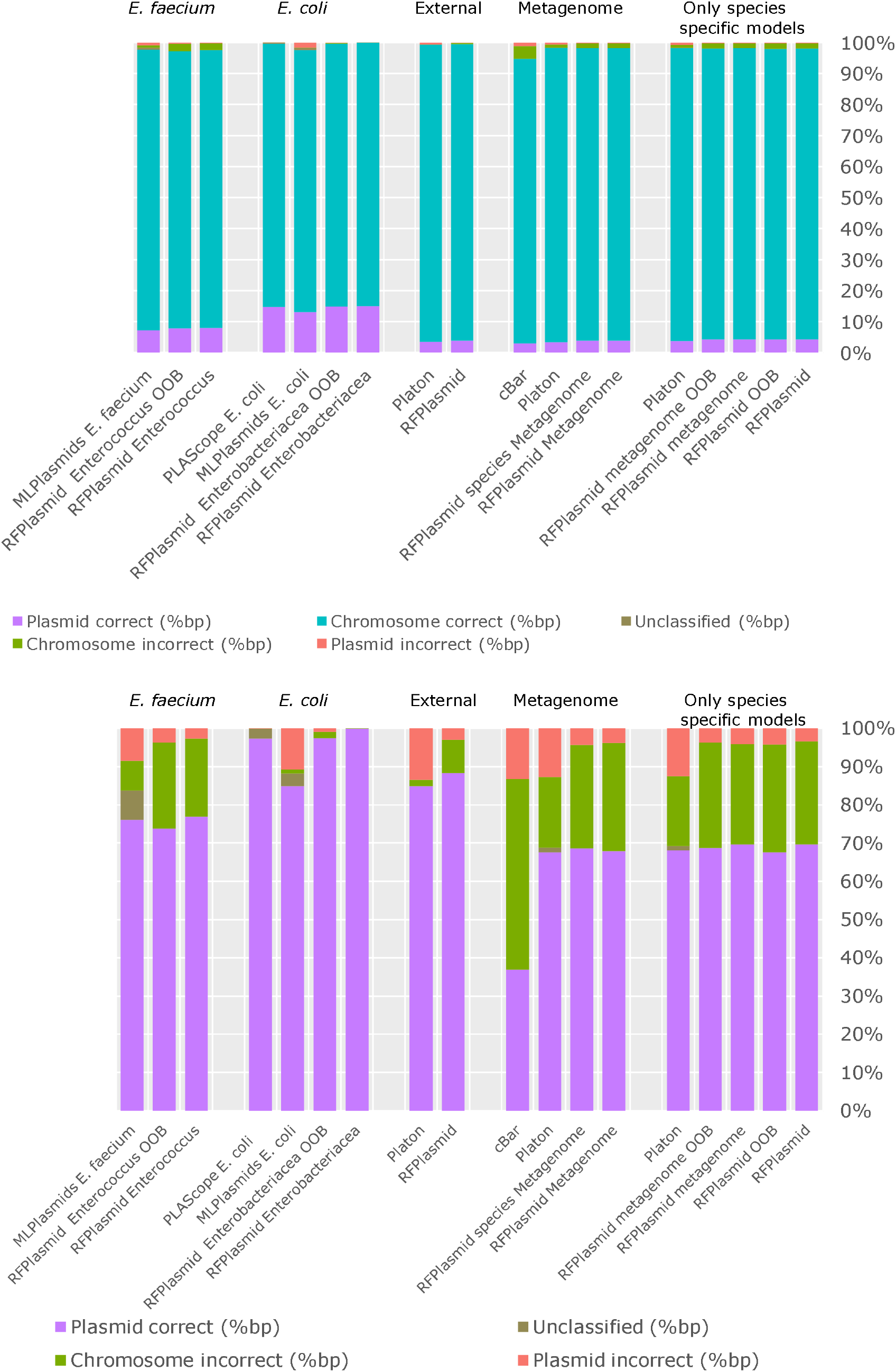
Comparison of RFPlasmid performance with existing tools. Shown are the OOB performance and prediction performance in percentages, calculated as basepairs predicted divided by the total basepairs for each plasmid correct, chromosome correct, chromosome incorrect and plasmid incorrect classified contig. (A) Chart including correctly identified chromosomal contigs and (B) Chart excluding correctly identified chromosomal contigs.

To investigate the performance of RFPlasmid on non-simulated data we also used 22 *E. coli* genomes, previously used by Schwenger *et al* (Schwengers et al., 2020). The error rate of RFPlasmid with non-simulated data is very low; only 0.52% of basepairs (85 contigs out of 2832 contigs) were incorrectly predicted with most of them (62 contigs out of 85 contigs) being small (<3kb) (Figure 6AB). Manual investigation of the larger incorrectly predicted contigs shows that 16 contigs contain phage encoding genes and three contigs a plasmid replication gene of which one encodes IncQ1, which is presumably integrated into the genome of isolate H69.

## Discussion and conclusion

We conclude that RFplasmid is able to predict chromosomal and plasmid contigs with error rates ranging from 0.002% to 4.66% (Figure 2A) and that the use of species specific models can be superior to a general plasmid prediction model. Single copy chromosomal marker genes, plasmid genes, k-mer content and length of contig all appear to be informative, however k-mer content is highly specific for species. Prediction of small contigs remains unreliable, since these contigs consists primarily of repeated sequences present in both plasmid and chromosome, e.g. transposases or because k-mer content or marker genes cannot be easily identified.

Comparison with existing methods shows that RFPlasmid generally performs equal or better to existing methods with a specificity and sensitivity up to 99%. RFPlasmid is the first described tool that can be used for 17 bacterial species and also includes a mode when the species is not in the database (e.g. also suitable for metagenomics assembly data). If a good reference set with well identified chromosomal and plasmid contigs of another bacterial species is available, an easy training option is implemented in RFPlasmid, to train a new model for this bacterial species. Our web-interface makes RFPlasmid accessible to the casual non-bioinformatician user, which will improve uptake of the use of our tool.

Improvements are still possible for RFPlasmid. Careful examination of the incorrectly classified contigs shows that these frequently contain many phage genes or transposases. Phages are often found on chromosomes, rarely on plasmids, therefore including a phage detection algorithm could certainly improve predictions, although that is out of scope for this study, as phage prediction has its own difficulties and complexities. Furthermore, phage-like plasmids have been detected (Galetti et al., 2019; Octavia et al., 2015) which would need to be investigated if it is possible to distinguish these from real phages. Smaller contigs which consisting solely of transposases (1-3 kb usually) are generally present on both chromosome and plasmid and these could be detected and marked as such. Integrated plasmids, such as the IncQ1 plasmid in the external dataset in *E. coli* isolate H69 show that some predictions will remain difficult. Other improvements could be the detection of rRNA operons, as these are usually chromosomally encoded or circularization detection for the detection of smaller plasmids (Schwengers et al., 2020). An evaluation of the combination of above mentioned features with species specific models would be interesting for future research

### Availability and requirements

Project name: RFPlasmid
Project home page: https://github.com/aldertzomer/RFPlasmid
Operating system(s): Linux (shell), platform independent
Programming language: Python, R, Bash
Other requirements: CheckM, Diamond
Optional: Jellyfish
License: e.g. AGPL probably.
Any restrictions to use by non-academics: None

## References

Alikhan, N. F., Zhou, Z., Sergeant, M. J., & Achtman, M. (2018). A genomic overview of the population structure of Salmonella. PLoS Genetics, 14(4), 1–13. https://doi.org/10.1371/journal.pgen.1007261

Antipov, D., Hartwick, N., Shen, M., Raiko, M., Lapidus, A., & Pevzner, P. A. (2016). PlasmidSPAdes: Assembling plasmids from whole genome sequencing data. Bioinformatics, 32(22), 3380–3387. https://doi.org/10.1093/bioinformatics/btw493

Arredondo-Alonso, S., Rogers, M. R. C., Braat, J. C., Verschuuren, T. D., Top, J., Corander, J., Willems, R. J. L., & Schürch, A. C. (2018). Mlplasmids: a User-Friendly Tool To Predict Plasmid- and Chromosome-Derived Sequences for Single Species. Microbial Genomics, 4(11). https://doi.org/10.1099/mgen.0.000224

Arredondo-Alonso, S., Willems, R. J., van Schaik, W., & Schürch, A. C. (2017). On the (im)possibility of reconstructing plasmids from whole-genome short-read sequencing data. Microbial Genomics, 3(10). https://doi.org/10.1099/mgen.0.000128

Bankevich, A., Nurk, S., Antipov, D., Gurevich, A. A., Dvorkin, M., Kulikov, A. S., Lesin, V. M., Nikolenko, S. I., Pham, S., Prjibelski, A. D., Pyshkin, A. V., Sirotkin, A. V., Vyahhi, N., Tesler, G., Alekseyev, M. A., & Pevzner, P. A. (2012). SPAdes: A new genome assembly algorithm and its applications to single-cell sequencing. Journal of Computational Biology, 19(5), 455–477. https://doi.org/10.1089/cmb.2012.0021

Buchfink, B., Xie, C., & Huson, D. (2015). Fast and sensitive protein alignment using DIAMOND. Nature Methods, 12(1), 59–60. https://doi.org/10.1038/nmeth.3176

Carattoli, A. (2009). Resistance plasmid families in Enterobacteriaceae. Antimicrobial Agents and Chemotherapy, 53(6), 2227–2238. https://doi.org/10.1128/AAC.01707-08

Carattoli, A., Zankari, E., Garciá-Fernández, A., Larsen, M. V., Lund, O., Villa, L., Aarestrup, F. M., & Hasman, H. (2014). In Silico detection and typing of plasmids using plasmidfinder and plasmid multilocus sequence typing. Antimicrobial Agents and Chemotherapy, 58(7), 3895–3903. https://doi.org/10.1128/AAC.02412-14

Dib, J. R., Wagenknecht, M., Farías, M. E., & Meinhardt, F. (2015). Strategies and approaches in plasmidome studies-uncovering plasmid diversity disregarding of linear elements? Frontiers in Microbiology, 5(MAY), 1–12. https://doi.org/10.3389/fmicb.2015.00463

Edgar, R. C. (2010). Search and clustering orders of magnitude faster than BLAST. Bioinformatics, 25(19), 2460–2461. https://doi.org/10.1093/bioinformatics/btq461

Galetti, R., Andrade, L. N., Varani, A. M., & Darini, A. L. C. (2019). A phage-like plasmid carrying blaKPC-2Gene in carbapenem-resistant pseudomonas aeruginosa. Frontiers in Microbiology, 10(MAR), 2–6. https://doi.org/10.3389/fmicb.2019.00572

Goessweiner-mohr, N., Arends, K., Keller, W., & Grohmann, E. (2014). Conjugation in Gram-Positive Bacteria. https://doi.org/10.1128/microbiolspec.PLAS-0004

Hamada, M., Ono, Y., Asai, K., Frith, M. C., & Hancock, J. (2017). Training alignment parameters for arbitrary sequencers with LAST-TRAIN. Bioinformatics, 33(6), 926–928. https://doi.org/10.1093/bioinformatics/btw742

Hyatt, D., Chen, G. L., LoCascio, P. F., Land, M. L., Larimer, F. W., & Hauser, L. J. (2010). Prodigal: Prokaryotic gene recognition and translation initiation site identification. BMC Bioinformatics, 11. https://doi.org/10.1186/1471-2105-11-119

Janitza, S., & Hornung, R. (2018). On the overestimation of random forest’s out-of-bag error. In PLoS ONE (Vol. 13, Issue 8). https://doi.org/10.1371/journal.pone.0201904

Johnson, T. J., & Nolan, L. K. (2009). Pathogenomics of the Virulence Plasmids of Escherichia coli. Microbiology and Molecular Biology Reviews, 73(4), 750–774. https://doi.org/10.1128/mmbr.00015-09

Jolley, K. A., Bray, J. E., & Maiden, M. C. J. (2018). Open-access bacterial population genomics: BIGSdb software, the PubMLST.org website and their applications [version 1; referees: 2 approved]. Wellcome Open Research, 3(0), 1–20. https://doi.org/10.12688/wellcomeopenres.14826.1

Lanza, V. F., de Toro, M., Garcillán-Barcia, M.P., Mora, A., Blanco, J., Coque, T. M., & de la Cruz, F. (2014). Plasmid Flux in Escherichia coli ST131 Sublineages, Analyzed by Plasmid Constellation Network (PLACNET), a New Method for Plasmid Reconstruction from Whole Genome Sequences. PLoS Genetics, 10(12). https://doi.org/10.1371/journal.pgen.1004766

Li, Y., Canchaya, C., Fang, F., Raftis, E., Ryan, K. A., Van Pijkeren, J. P., Van Sinderen, D., & O’Toole, P. W. (2007). Distribution of megaplasmids in Lactobacillus salivarius and other Lactobacilli. Journal of Bacteriology, 189(17), 6128–6139. https://doi.org/10.1128/JB.00447-07

Liaw, A., & Wiener, M. (2002). Classification and Regression by randomForest. R News, 2(3), 18–22.

Marçais, G., & Kingsford, C. (2011). A fast, lock-free approach for efficient parallel counting of occurrences of k-mers. Bioinformatics, 27(6), 764–770. https://doi.org/10.1093/bioinformatics/btr011

Octavia, S., Sara, J., & Lan, R. (2015). Characterization of a large novel phage-like plasmid in Salmonella enterica serovar Typhimurium. FEMS Microbiology Letters, 362(8), 1–9. https://doi.org/10.1093/femsle/fnv044

Parks, D. H., Imelfort, M., Skennerton, C. T., Hugenholtz, P., & Tyson, G. W. (2015). CheckM: Assessing the quality of microbial genomes recovered from isolates, single cells, and metagenomes. Genome Research, 25(7), 1043–1055. https://doi.org/10.1101/gr.186072.114

Reis-Cunha, J. L., Bartholomeu, D. C., Manson, A. L., Earl, A. M., & Cerqueira, G. C. (2019). ProphET, prophage estimation tool: A standalone prophage sequence prediction tool with self-updating reference database. PLoS ONE, 14(10), 1–9. https://doi.org/10.1371/journal.pone.0223364

Robertson, J., & Nash, J. H. E. (2018). MOB-suite: software tools for clustering, reconstruction and typing of plasmids from draft assemblies. Microbial Genomics, 4(8). https://doi.org/10.1099/mgen.0.000206

Royer, G., Decousser, J. W., Branger, C., Dubois, M., Médigue, C., Denamur, E., & Vallenet, D. (2018). PlaScope: A targeted approach to assess the plasmidome from genome assemblies at the species level. Microbial Genomics, 4(9), 1–8. https://doi.org/10.1099/mgen.0.000211

Rozov, R., Kav, A. B., Bogumil, D., Shterzer, N., Halperin, E., Mizrahi, I., & Shamir, R. (2017). Recycler: An algorithm for detecting plasmids from de novo assembly graphs. Bioinformatics, 33(4), 475–482. https://doi.org/10.1093/bioinformatics/btw651

Rozwandowicz, M., Brouwer, M. S. M., Fischer, J., Wagenaar, J. A., Gonzalez-Zorn, B., Guerra, B., Mevius, D. J., & Hordijk, J. (2018). Plasmids carrying antimicrobial resistance genes in Enterobacteriaceae. Journal of Antimicrobial Chemotherapy, 73(5), 1121–1137. https://doi.org/10.1093/jac/dkx488

Schwengers, O., Barth, P., Falgenhauer, L., Hain, T., Chakraborty, T., & Goesmann, A. (2020). Platon: identification and characterization of bacterial plasmid contigs in short-read draft assemblies exploiting protein-sequence-based replicon distribution scores. BioRxiv, 2020.04.21.053082. https://doi.org/10.1101/2020.04.21.053082

Sengupta, M., & Austin, S. (2011). Prevalence and significance of plasmid maintenance functions in the virulence plasmids of pathogenic bacteria. Infection and Immunity, 79(7), 2502–2509. https://doi.org/10.1128/IAI.00127-11

Smillie, C., Garcillán-Barcia, M. P., Francia, M. V., Rocha, E. P. C., & de la Cruz, F. (2010). Mobility of Plasmids. Microbiology and Molecular Biology Reviews, 74(3), 434–452. https://doi.org/10.1128/mmbr.00020-10

Wattam, A. R., Davis, J. J., Assaf, R., Boisvert, S., Brettin, T., Bun, C., Conrad, N., Dietrich, E. M., Disz, T., Gabbard, J. L., Gerdes, S., Henry, C. S., Kenyon, R. W., Machi, D., Mao, C., Nordberg, E. K., Olsen, G. J., Murphy-Olson, D. E., Olson, R., … Stevens, R. L. (2017). Improvements to PATRIC, the all-bacterial bioinformatics database and analysis resource center. Nucleic Acids Research, 45(D1), D535–D542. https://doi.org/10.1093/nar/gkw1017

Wick, R. R., Judd, L. M., Gorrie, C. L., & Holt, K. E. (2017). Unicycler: Resolving bacterial genome assemblies from short and long sequencing reads. PLoS Computational Biology,13(6), 1–22. https://doi.org/10.1371/journal.pcbi.1005595

Wick, R. R., Schultz, M. B., Zobel, J., & Holt, K. E. (2015). Bandage: Interactive visualization of de novo genome assemblies. Bioinformatics, 31(20), 3350–3352. https://doi.org/10.1093/bioinformatics/btv383

Xie, Z., & Tang, H. (2017). ISEScan: automated identification of insertion sequence elements in prokaryotic genomes. Bioinformatics (Oxford, England), 33(21), 3340–3347. https://doi.org/10.1093/bioinformatics/btx433

Zankari, E., Hasman, H., Cosentino, S., Vestergaard, M., Rasmussen, S., Lund, O., Aarestrup, F. M., & Larsen, M. V. (2012). Identification of acquired antimicrobial resistance genes. Journal of Antimicrobial Chemotherapy, 67(11), 2640–2644. https://doi.org/10.1093/jac/dks261

Zhou, F., & Xu, Y. (2010). cBar: A computer program to distinguish plasmid-derived from chromosome-derived sequence fragments in metagenomics data. Bioinformatics, 26(16), 2051–2052. https://doi.org/10.1093/bioinformatics/btq299

